# Transmission of highly virulent CXCR4 tropic HIV-1 through the mucosal route in an individual with a wild-type CCR5 genotype

**DOI:** 10.1101/2023.09.15.557832

**Authors:** Manukumar Honnayakanahalli Marichannegowda, Saini Setua, Meera Bose, Eric Sanders-Buell, David King, Michelle Zemil, Lindsay Wieczorek, Felisa Diaz-Mendez, Nicolas Chomont, Rasmi Thomas, Leilani Francisco, Leigh Anne Eller, Victoria R. Polonis, Sodsai Tovanabutra, Yutaka Tagaya, Nelson L. Michael, Merlin L. Robb, Hongshuo Song

## Abstract

Nearly all transmitted/founder (T/F) HIV-1 are CCR5 (R5)-tropic. While previous evidence suggested that CXCR4 (X4)-tropic HIV-1 are transmissible, detection was not at the earliest stages of acute infection. Here, we identified an X4-tropic T/F HIV-1 in a participant in acute infection cohort. Coreceptor assays demonstrated that this T/F virus is strictly CXCR4 tropic. The participant experienced significantly faster CD4 depletion compared with R5 virus infected participants in the same cohort. Naïve and central memory CD4 subsets declined faster than effector and transitional memory subsets. All CD4 subsets, including naïve, were productively infected. Increased CD4^+^ T cell activation was observed over time. This X4-tropic T/F virus is resistant to broadly neutralizing antibodies (bNAbs) targeting V1/V2 and V3 regions. These findings demonstrate that X4-tropic HIV-1 is transmissible through the mucosal route in people with the wild-type CCR5 genotype and have implications for understanding the transmissibility and immunopathogenesis of X4-tropic HIV-1.

## Introduction

The vast majority of T/F HIV-1 are CCR5 tropic^1,2^. While the biological mechanisms underlying the transmission advantage of R5 tropic HIV-1 remain incompletely understood, previous evidence suggested that X4 tropic HIV-1 are transmissible. The best evidence comes from the identification of X4 viruses in people homozygous for CCR5 delta 32 allele^3–5^. These studies suggested that clinical HIV-1 infection could be established and maintained by the CXCR4 coreceptor. Moreover, the identification of X4 viruses in recently diagnosed HIV-1 infections also indicated that X4 viruses are transmissible, although the chance of transmission is low^6,7^. However, one caveat of the previous studies is that viral identification was not at the earliest stage of acute HIV infection. Therefore, the exact phenotype of the transmitted virus was unable to be characterized. Because HIV-1 could also use alternative coreceptors other than CCR5 and CXCR4, as assessed by *in vitro* assays, to establish clinical infection^5,8^, the possibility that viral transmission in people homozygous for CCR5 delta 32 was mediated by alternative coreceptors could not be completely excluded. To our knowledge, a T/F HIV-1 exclusively using CXCR4 has not been identified so far and it remains unclear whether X4 tropic HIV-1 without capacity to use CCR5 can be transmitted through the mucosal route in people with a wild-type CCR5 genotype.

We identified a participant (40700) who was infected by an X4 tropic T/F HIV-1 in the RV217 Thailand cohort, a longitudinal study of individuals at high risk of HIV-1 infection who were followed while still uninfected with twice weekly HIV-1 RNA testing^9^. The risk factor for HIV-1 transmission was men who have sex with men (MSM). Participant 40700 had a wild-type CCR5 genotype and expressed normal levels of CCR5 on CD4^+^ T cells. An unusually fast CD4 depletion was observed in participant 40700 upon HIV-1 transmission. We comprehensively characterized the genetic and phenotypic property related to the immunopathogenesis of this highly pathogenic T/F virus.

## Results

### Identification and characterization of the X4 tropic T/F virus in participant 40700

Participant 40700 was identified in the RV217 Thailand cohort and was infected by a CRF01_AE HIV-1. The first large volume blood draw was 15 days from the first HIV-1 RNA-positive test and 23 days from the last HIV-1 RNA-negative test. Analysis of near full-length viral genomes obtained by single genome amplification (SGA) identified a homogenous viral lineage at day 15 (Fig. 1a). Genetic analysis using the Poisson-Fitter tool^10^ showed that the frequency distribution of the Hamming Distance followed Poisson distribution and the sequences exhibited a star-like phylogeny (Fig. 1b). The estimated days from HIV-1 transmission (based on the Poisson-Fitter tool) was 26. These results demonstrated that participant 40700 was identified during acute HIV infection (AHI) which was established by a single T/F virus.

**Fig. 1.**
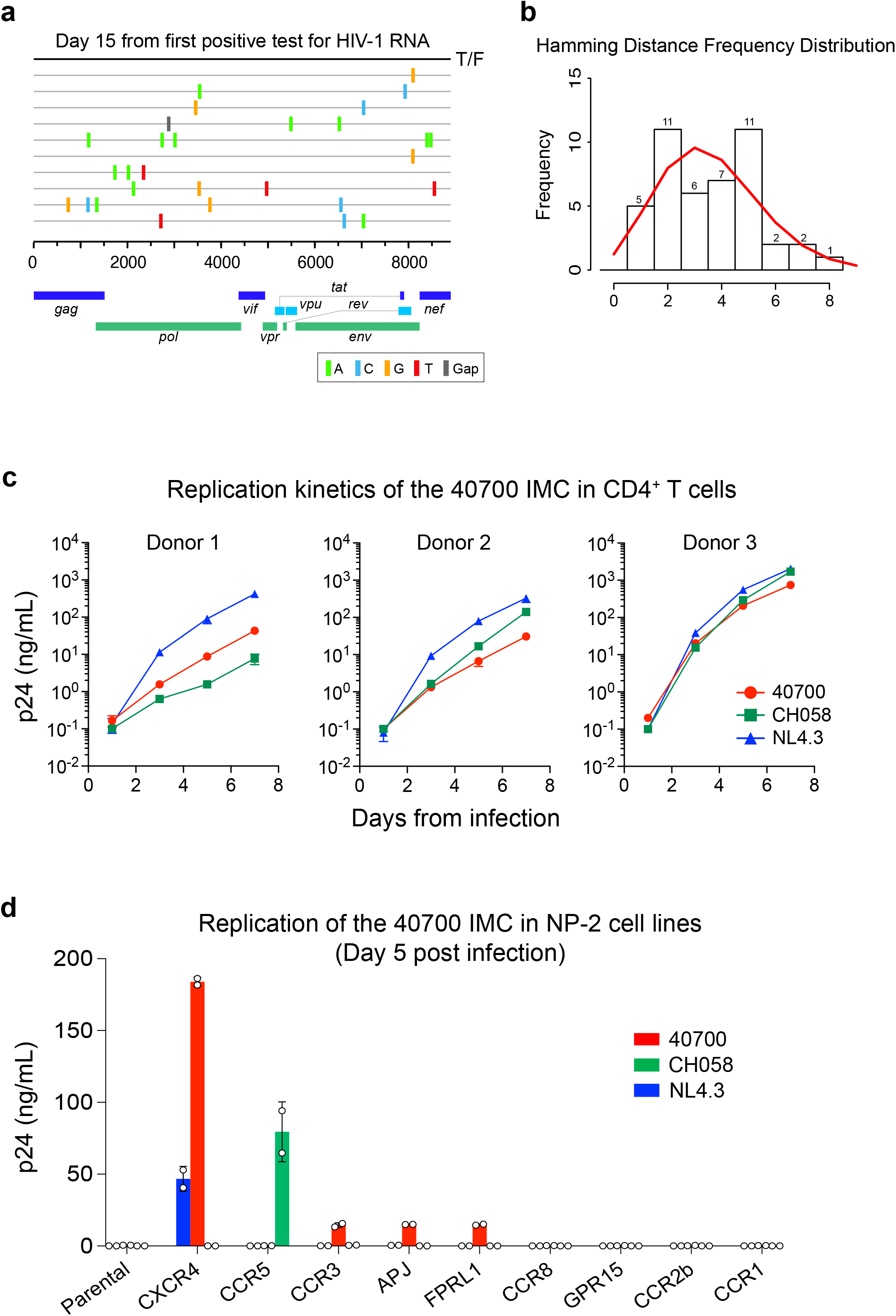
Genetic and phenotypic characterization of the 40700 T/F virus. **a,** Highlighter plot showing near-full length viral genomes obtained at day 15 from the 1^st^ positive test for HIV-1 RNA. The sequences were obtained by SGA. The consensus sequence (black line on the top) is used as the master sequence. Genetic substitutions compared to the consensus sequence are color coded. **b,** Frequency distribution of the pair-wise Hamming Distance of the day 15 viral sequences. The frequency distribution follows Poisson distribution as determined by the Poisson-Fitter tool in the Los Alamos HIV Sequence Database**. c**, Replication kinetics of the 40700 IMC in purified CD4^+^ T cells from three healthy donors. The X4 tropic strain NL4.3 and an R5 tropic T/F virus CH058 were used as controls. All infections were performed in triplicate. The error bar represents the standard deviation (SD). **d**, Determination of coreceptor usage of the 40700 IMC in a panel of NP-2 cell lines expressing different HIV-1 coreceptors. The p24 concentration in the culture supernatant was determined on day 5 post infection. The X4 tropic strain NL4.3 and an R5 tropic T/F virus CH058 were used as controls. The infections were performed in duplicate. The error bar shows the SD.

Coreceptor prediction using Geno2Pheno showed a high likelihood of CXCR4 usage of the 40700 T/F virus (FPR = 0.1%)^11^. This observation prompted us to phenotypically characterize the coreceptor usage of this T/F virus. Coreceptor inhibition assay in TZM-bl cell line showed that the entry activity of the 40700 T/F pseudovirus can be completely blocked by the CXCR4 inhibitor AMD3100 while the CCR5 inhibitor Maraviroc showed no inhibition at the highest concentration (100 µM) (Extended Data Fig. 1a). Coreceptor assay in NP-2 cell lines demonstrated that the 40700 T/F pseudovirus can infect the NP-2 CXCR4 cell line with high efficiency while no infectivity was observed in the NP-2 CCR5 cell line (Extended Data Fig. 1b). These data suggested that the 40700 T/F virus uses CXCR4 exclusively.

To better characterize the phenotype of the 40700 T/F virus, we constructed a full-length infectious molecular clone (IMC) (Extended Data Fig. 2). A viral replication assay in purified CD4^+^ T cells from three healthy donors showed that the 40700 T/F IMC was replication competent in primary CD4^+^ T cells (Fig. 1c). Overall, its replication kinetics was comparable to the X4 tropic strain NL4.3 and an R5-tropic T/F virus CH058 (Fig. 1c). We then determined the coreceptor usage of the 40700 IMC in a panel of NP-2 cell lines expressing CCR5, CXCR4, as well as another seven alternative coreceptors (Fig. 1d). Consistent with the results of the pseudovirus, the 40700 IMC replicated in the NP-2 CXCR4 cell line with high efficiency. No viral replication was detected in the NP-2 CCR5 cell line (Fig. 1d). The 40700 IMC could also infect CCR3, APJ and FPRL1 cell lines with low efficiency (Fig. 1d).

A CCR5 genotyping showed that participant 40700 did not have the CCR5 delta32 mutation (Methods). We analyzed the CCR5 and CXCR4 expression on CD4^+^ T cells of 40700 isolated from day 20 (from the first positive test for HIV-1 RNA) (Extended Data Fig. 3). The CCR5 and CXCR4 expression in 40700 showed similar pattern as previously observed in two participants infected by R5 viruses in the same cohort^12^. The CCR5 expression was high on the effector memory (EM) and transitional memory (TM) CD4 subsets, low on the central memory (CM) subset, and was undetectable on the naïve subset. The CXCR4 expression showed a reciprocal pattern (Extended Data Fig. 3). Taken together, these data demonstrated that participant 40700 was infected by a CXCR4 tropic T/F virus lacking demonstrable CCR5 tropism, and the transmission of X4 virus in 40700 could not be explained by unusual expression of the CCR5 and CXCR4 coreceptors.

### Rapid CD4^+^ T cell depletion in participant 40700

Participant 40700 experienced significantly faster CD4^+^ T cell decline compared to infections established by R5 virus in the RV217 Thailand cohort (Fig. 2a). The CD4 count dropped from 707 cells/µL to 130 cells/µL during the first 223 days of infection before ART initiation (40700 was on and off ART since day 266 as described in Methods) (Fig. 2b). The rate of CD4 decline was 1.6 cells/µL/day in 40700, while the median rate of CD4 decline in the R5 group was 0.27 cells/µL/day (Fig. 2a). By analyzing the mean and variance of the participant specific CD4 slopes with a Linear Mixed Effect (LME) model, we were able to confirm that 40700 had a statistically significant rate of CD4^+^ T cell decline compared with the R5 group (*P* < 0.001; normal distribution test) (Fig. 2a). The CD8 T cell count was relatively stabilized after AHI and the plasma viral load (VL) was sustained above 10^5^ copies/mL (Fig. 2b). Analysis of the dynamics of each CD4 subset showed that the naïve subset declined faster (0.97 cells/µL/day) than the memory subsets (Fig. 2c). Among the memory CD4 subsets, the central memory (CM) subset declined faster (0.42 cells/µL/day) than effector memory (EM) and transitional memory (TM) subsets (0.17 and 0.07 cells/µL/day, respectively) (Fig. 2c). These data suggested that the rapid CD4 depletion in 40700 was mainly due to the loss of naïve and CM CD4^+^ T cells.

**Fig. 2.**
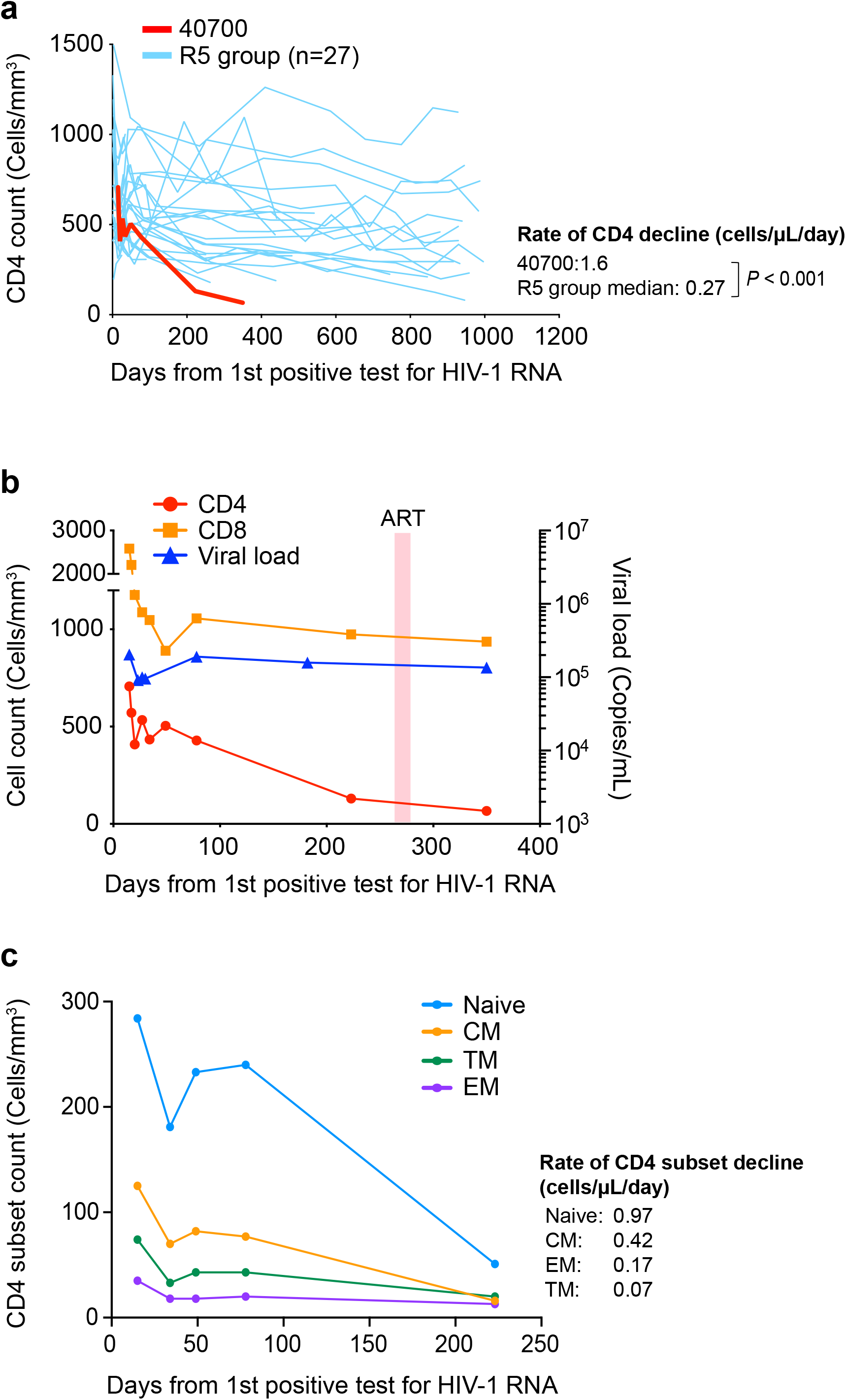
Rapid CD4^+^ T cell depletion in participant 40700 after HIV-1 transmission. **a**, Dynamics of CD4^+^ T cells in participant 40700 compared to R5 HIV-1 infected participants in the RV217 Thailand cohort. The rate of CD4 decline was calculated using a linear mixed effect model (LME). The statistical significance was determined using a normal distribution test. **b**, Dynamics of CD4 count, CD8 count, and plasma VL in participant 40700. The time frame when participant 40700 was on ART is highlighted in red (day 266 to 282). **c**, Dynamics of each CD4 subset in participant 40700 before ART initiation. Four different CD4 subsets (naïve, CM, EM, and TM) are color coded. The rate of CD4 subset decline was calculated by linear regression.

### Broad CD4 subset targeting of the 40700 T/F virus

CXCR4 tropic HIV-1 are considered to have a broader CD4 subset targeting than R5 viruses^13^. In X4-tropic SHIV infected macaques, the resting naïve CD4^+^ T cells were productively infected and rapidly depleted^14^. The strict X4 tropism of the 40700 T/F virus provides a unique opportunity to determine the CD4 subset preference and pathogenesis of X4 virus in natural HIV-1 infection. The faster depletion of the naïve and CM CD4 subsets in 40700 indicated that they could be preferentially infected due to relatively high levels of CXCR4 expression (Extended Data Fig. 3). Quantification of total and integrated HIV-1 DNA at day 78 showed that both the total and integrated HIV-1 DNA were detectable in all CD4 subsets (Fig. 3a). The CM CD4 subset contained the highest level of both the total and integrated HIV-1 DNA (Fig. 3a). Possibly due to the relatively low level of CXCR4 expression, the EM and TM subsets contained lower amount of total HIV-1 DNA compared with the CM and naïve subsets (Fig. 3a). In the naïve subset, the level of total HIV-1 DNA was 9.1-fold higher than the integrated DNA, while in the memory subsets, the total HIV-1 DNA was approximately 2-fold higher than the integrated DNA (Fig. 3a). This observation indicated that while the viruses could enter the naïve CD4^+^ T cells with high efficiency, the integration process could be defective or delayed due to the resting status of the naïve cells.

**Fig. 3.**
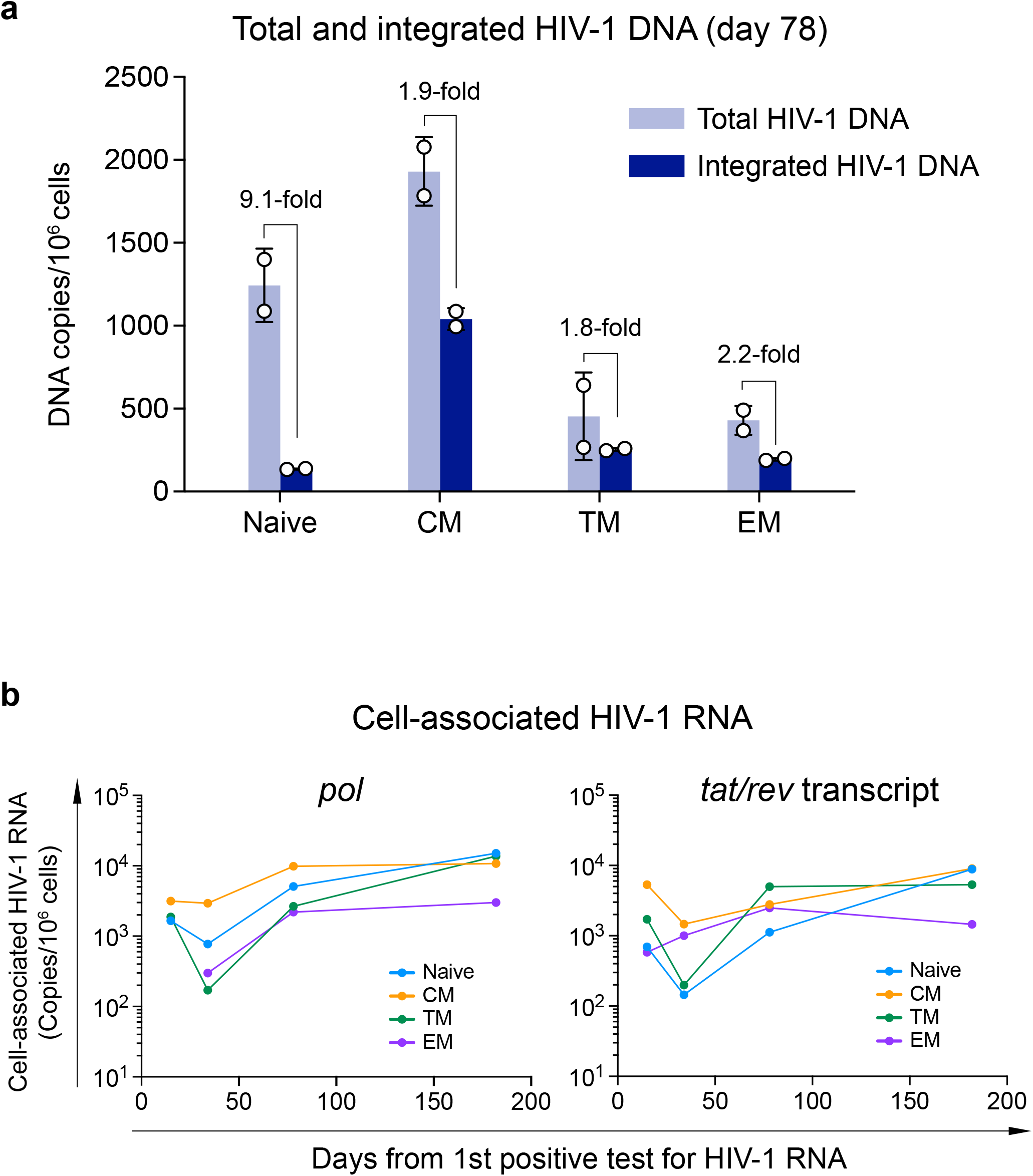
Quantification of total and integrated HIV-1 DNA and cell-associated HIV-1 RNA in participant 40700. **a**, Total (light blue) and integrated (dark blue) HIV-1 DNA was quantified in each sorted CD4 subset at day 78 (from the 1^st^ positive test for HIV-1 RNA). The results are shown as DNA copies/10^6^ cells. The experiments were performed in duplicate. The error bar shows the SD. **b**, Dynamics of cell-associated HIV-1 RNA in each CD4 subset. The cell-associated HIV-1 RNA in each CD4 subset was quantified by amplifying the *pol* region and the *tat/rev* transcript. Different CD4 subsets are color coded. The *pol* region in the EM CD4 subset was not amplifiable at the first time point (day 20). The cell-associated HIV-1 RNA viral load is shown as copies/10^6^ cells.

To determine which CD4 subsets were productively infected in 40700, we quantified the cell-associated HIV-1 RNA in longitudinal samples by amplifying both the *pol* region and the *tat/rev* transcript (Fig. 3b). Because the *tat/rev* transcript was generated at relatively late stage of virus life cycle, it could more accurately reflect the productive infection status. The results showed that all CD4 subsets were productively infected (Fig. 3b). In contrast, in RV217 Thailand participants who only harbored R5 viruses, the cell-associated HIV-1 RNA were undetectable in naïve CD4 subset as demonstrated in our recent study^12^. Quantification of the *pol* region and the *tat/rev* transcript showed similar dynamics (Fig. 3b). At earlier time points, the CM CD4 subset contained relatively high level of cell-associated HIV-1 RNA (Fig. 3b). The naïve and TM subsets contained relatively low level of cell-associated RNA initially but achieved similar level as the CM subset at the last time point. The EM subset had the lowest level of cell-associated RNA at the last time point (Fig. 3b). These results suggested that the CM CD4 subset was preferentially targeted by the 40700 T/F virus during early infection.

### Heightened CD4^+^ T cell activation over time

Immune activation is correlated with CD4 decline in HIV-1 infection^15,16^. The rapid CD4 depletion in 40700 prompted us to determine the level of CD4^+^ T cell activation over time. Quantification of the expression of T cell activation markers showed that after acute infection, coincident with a brief drop of plasma VL (Fig. 2b), there was a temporary decrease of CD4 activation and proliferation (Fig. 4). However, instead of achieving a steady-state as observed in most HIV infected people^15^, the level of CD4 activation increased after this brief decline. Among different CD4 subsets, the TM subset expressed the highest level of activation and proliferation markers (Fig. 4). The naïve CD4 subset was negative for all investigated markers except for CD38 (Fig. 4). The lack of Ki-67 expression in the naïve subset suggested that as previously observed in X4-tropic SHIV infected macaques^14^, the naïve CD4^+^ T cells which were productively infected and depleted in 40700 were largely quiescent. The level of CD4 activation and proliferation decreased upon ART. However, the expression of PD-1 did not decrease immediately after treatment (Fig. 4).

**Fig. 4.**
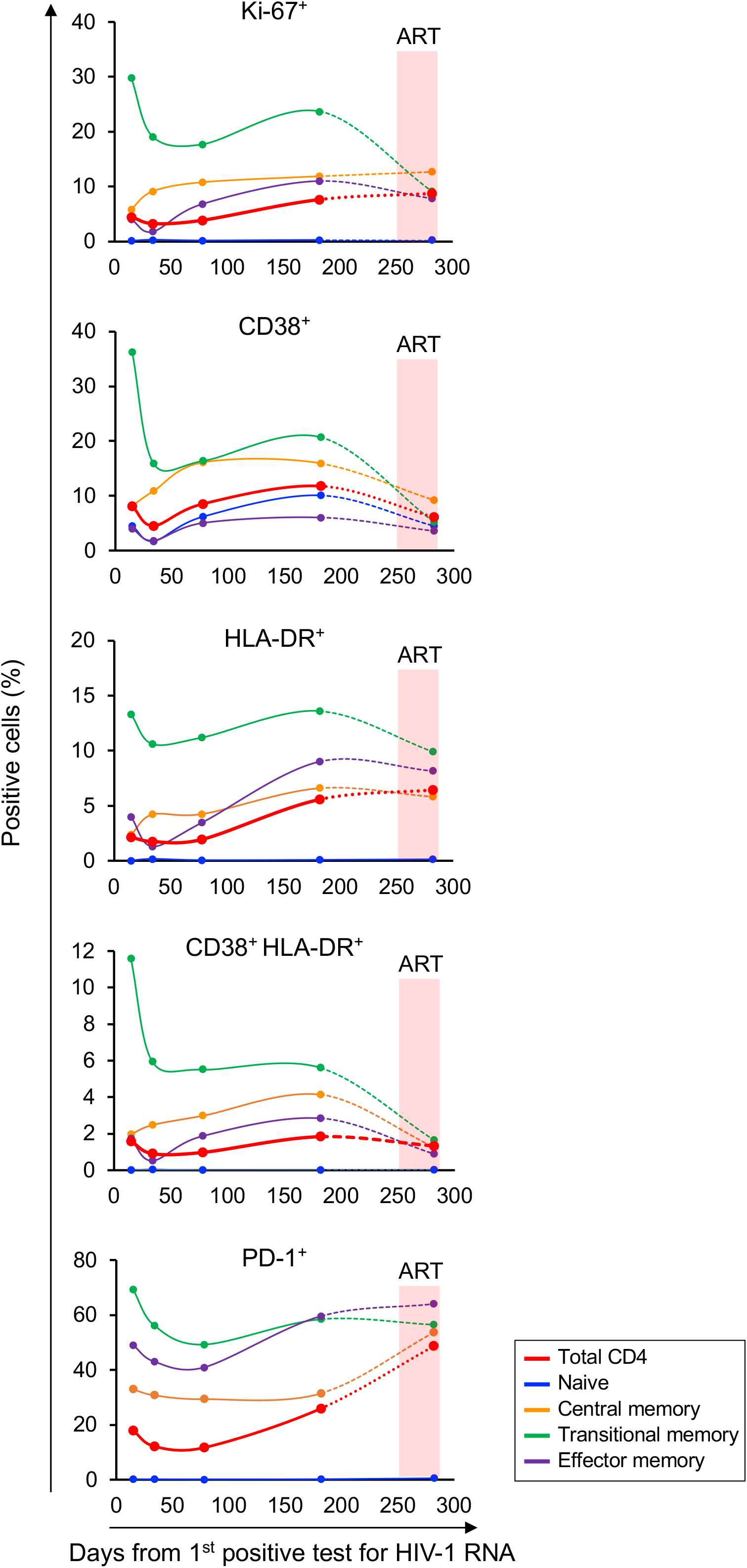
Determination of CD4^+^ T cell activation and proliferation in participant 40700. The expression of activation and proliferation markers on total CD4^+^ T cells as well as on each CD4 subset was determined by flowcytometry. The percentage of cells positive for each marker is shown. The time frame when participant 40700 was on ART (from day 266 to day 282) is highlighted in red.

### The 40700 T/F virus is resistant to bNAbs targeting the V1/V2 and V3 regions

X4 tropic HIV-1 from different subtypes tend to be more resistant to broadly neutralizing antibodies (bNAbs) targeting the V3 region^12,17–19^. Determination of neutralization susceptibility to a panel of bNAbs showed that while the 40700 T/F virus can be neutralized by bNAbs targeting the CD4 binding site and MPER, it was completely resistant to all investigated bNAbs targeting the V1/V2 and V3 regions (Fig. 5a). Investigation of the kinetics of autologous neutralization showed that participant 40700 developed relatively low level of autologous neutralization activity compared with two infections established by R5 T/F viruses (Fig. 5b).

**Fig. 5.**
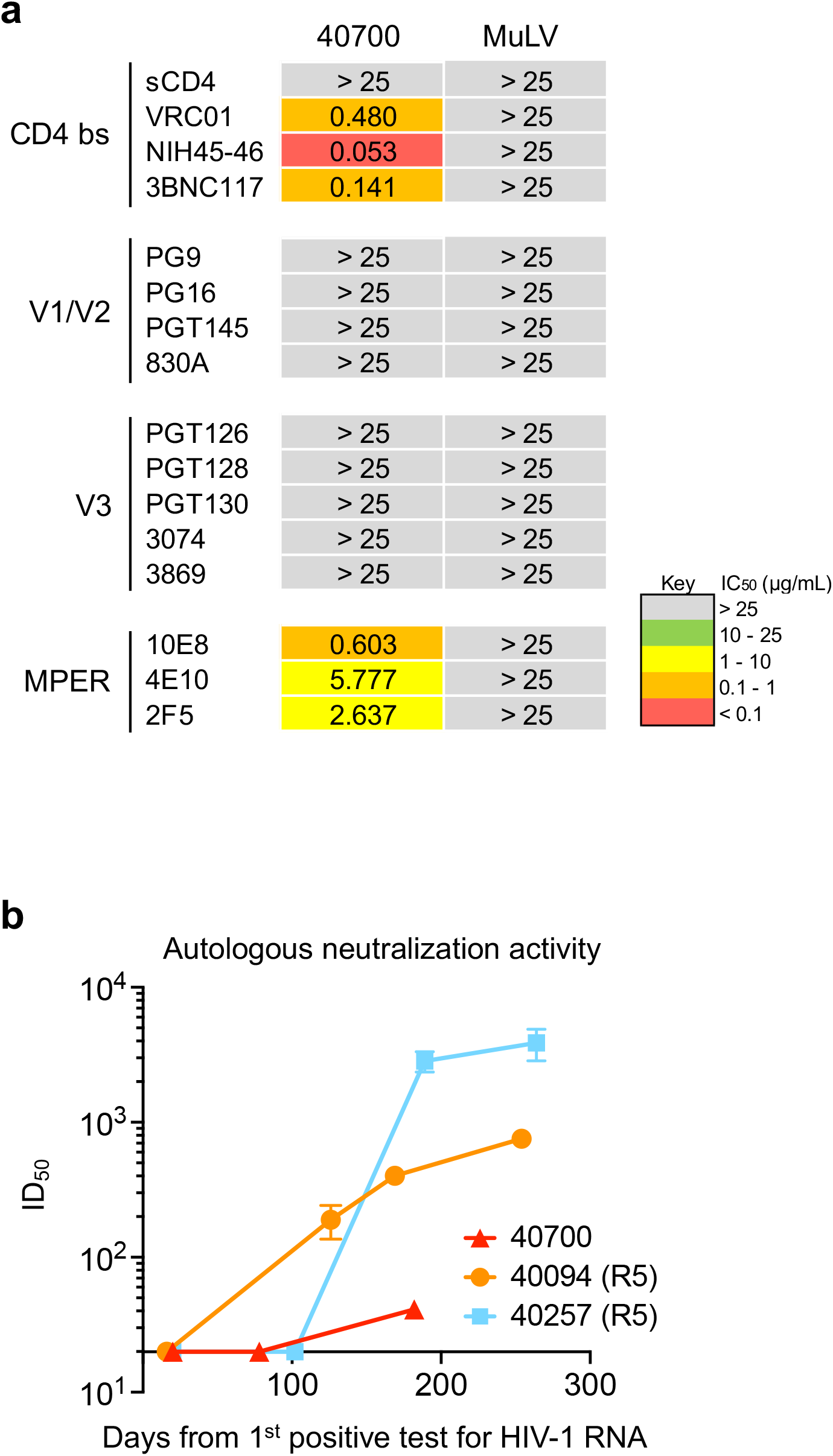
Neutralization susceptibility of the 40700 T/F virus to a panel of bNAbs and the autologous neutralization activity induced in 40700. **a,** Susceptibility of the 40700 T/F virus to a panel of bNAbs targeting different regions of the HIV-1 envelope. The IC_50_ (µg/mL) of each bNAb is shown. The murine leukemia virus (MuLV) was used as negative control. **b,** Autologous neutralization activity induced by the 40700 T/F virus. Two participants infected by R5 tropic T/F viruses (40257 and 40094) were used as controls.

Analysis of V3 amino acid sequence identified that the conserved V3 N301 glycan site is lost in 40700 due to the T303I substitution (Extended Data Fig. 4a). Indeed, substitutions at the N301 glycan exist in nearly all X4 viruses identified in the RV217 Thailand cohort^12^. As most of the CRF01_AE viruses, the N332 glycan site shifted to the N334 position in 40700 (Extended Data Fig. 4a). In comparison to the CRF01_AE consensus sequence, the 40700 T/F virus has two positively charged amino acid substitutions in V3, including the Q313R mutation at the V3 crown (Extended Data Fig. 4a). Moreover, the 40700 T/F virus has a longer V2 loop and an additional glycan stie in V2 region compared with most R5 T/F viruses in the same cohort (Extended Data Fig. 4b-c). These findings identified unique genetic feature and glycan arrangement in V1/V2 and V3 regions of the 40700 T/F virus.

Investigation of longitudinal envelope evolution in 40700 showed that the viral population diversified from the T/F virus by accumulation of point mutations (Fig. 6). Interestingly, during early viral evolution, the variable loops were relatively conserved while most of the fixed or predominant mutations emerged in the conserved regions (Fig. 6). This could be associated with relatively low level of autologous neutralization response in 40700.

**Fig. 6.**
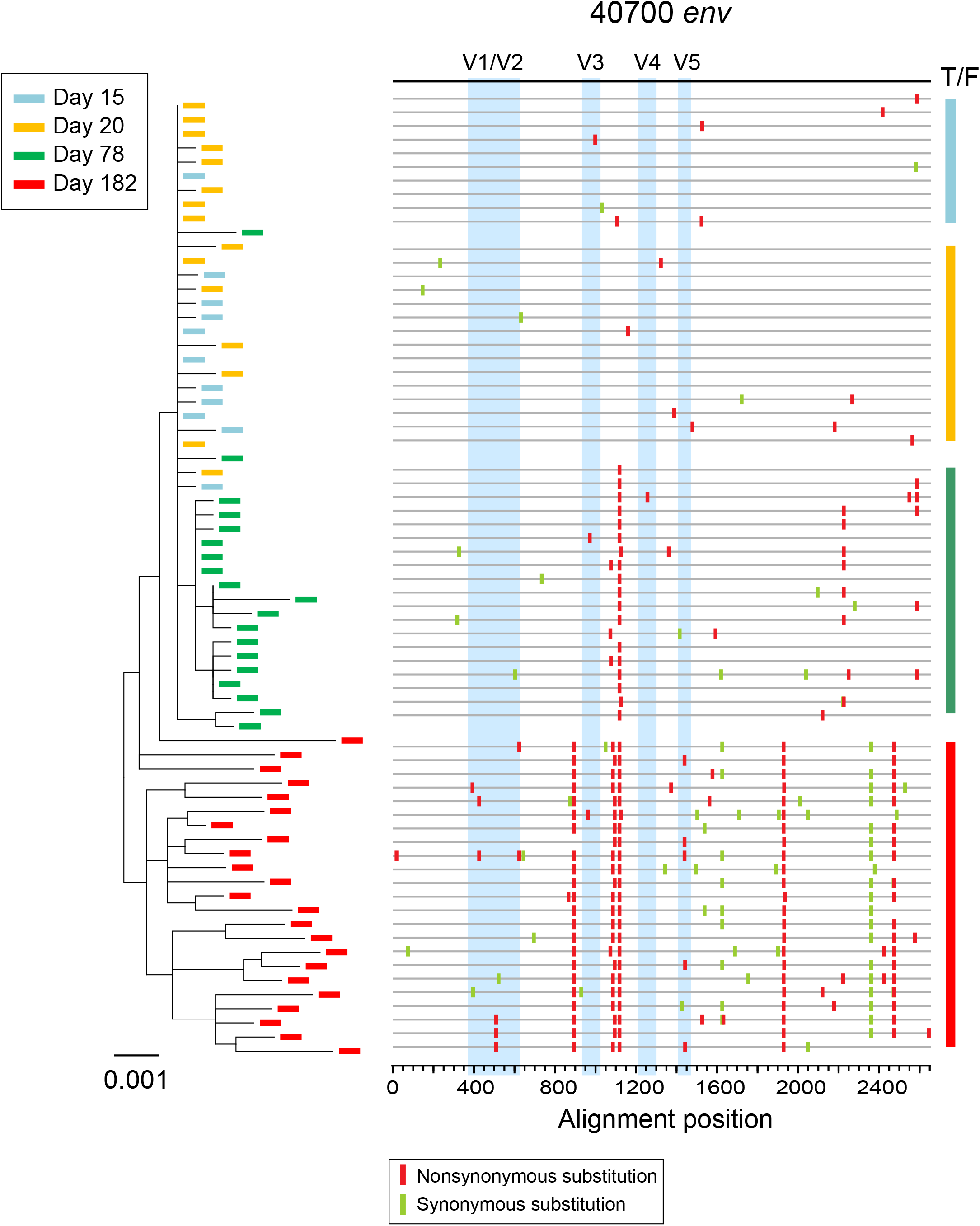
Phylogenetic tree and highlighter plot showing longitudinal viral evolution in participant 40700. Longitudinal *env* sequences were obtained by SGA. Evolution of the 40700 T/F virus is illustrated by phylogenetic tree (left) and highlighter plot (right). Sequences from different time points (days from the 1^st^ positive test for HIV-1 RNA) are color coded. The phylogenetic tree was constructed using the maximum likelihood method. In the highlighter plot, the black line on the top represents the T/F virus. The red and green tics indicate non-synonymous and synonymous substitutions compared to the T/F virus, respectively. The variable loops are shaded in blue.

## Discussion

The current study demonstrates that HIV-1 with strict X4 tropism can be transmitted through the mucosal route in a person with wild-type CCR5 genotype and can cause rapid CD4 depletion. This finding, together with the fact that the 40700 T/F virus is resistant to bNAbs targeting the V1/V2 and V3 regions, necessitate the need to monitor the transmission and spread of highly pathogenic X4 HIV-1 which tend to be resistant to bNAbs targeting the variable loops^12,17–19^.

While it has been well established that R5 tropic HIV-1 have transmission advantage, the underlying mechanisms remain poorly understood. A long-term question is why X4 viruses are less transmissible while CXCR4 is expressed on a broader range of CD4^+^ T cells than CCR5. The current study, together with our recent findings^12^, provide evidence that the different CD4 subset preference could be an important determinant underlying the distinct transmissibility of R5 and X4 HIV-1. In our recent study, tracking coreceptor switch of the T/F virus demonstrated that upon the origin of the earliest X4 virus, the viral population diverged *in vivo* in terms of CD4 subset targeting^12^. While the R5 population remained predominant in the EM and TM CD4 subsets, the emerging X4 viruses lost advantage in the EM and TM subsets while gained advantage in the CM and naïve subsets. In participant 40700, the CM CD4 subset was preferentially infected during early infection while the EM subset was less infected by this X4 tropic T/F virus. Because the EM CD4^+^ T cells are more abundant in mucosal tissues than the CM and naïve CD4^+^ T cells^20–22^, the replication advantage of R5 virus in the EM CD4^+^ T cells could provide an advantage of mucosal transmission. Moreover, the EM and TM cells are likely to have higher viral burst size than the naïve and CM cells due to their more activated status, and thus release more virions into the plasma. Indeed, the R5 viruses remained predominant in plasma in all participants harboring X4 variants in the RV217 Thailand cohort^12^. This could be another reason for the transmission advantage of R5 tropic HIV-1. Our findings also indicate that the EM and TM CD4 subsets could play a more important role in mediating mucosal HIV-1 transmission than CM and naïve subsets (otherwise, the X4 virus would have a higher chance to be transmitted given their advantage in the CM and naïve subsets). A better understanding of the CD4 subset preference in determining the transmissibility of HIV-1 could open new possibilities for preventing HIV-1 transmission (for example, by downregulation of CCR5 expression on the EM and TM CD4 subsets).

While the altered CD4 subset preference towards the naïve and CM cells may compromise the transmissibility of X4 tropic HIV-1, it could enhance the pathogenicity. In 40700, the rapid CD4 depletion was mainly due to the loss of naïve and CM CD4 subsets. In line with this finding, our recent study on coreceptor switch showed that upon the origin of the X4 viruses, the CM and naïve CD4 subsets declined faster than the EM and TM subsets^12^. Similar as previously observed in macaques infected by X4-tropic SHIV^14^, the resting naïve CD4^+^ T cells were productively infected and depleted in 40700. The exponentially increased viral load in the naïve subset was also similar as previously observed in the macaque model^14^. The mechanisms for productive infection and depletion of the resting naïve CD4^+^ T cells *in vivo* require further investigation. In addition to direct infection of the naïve CD4^+^ T cells, the high viral burden in the CM CD4 subset, which is essential for CD4 hemostasis as shown in SIV infected natural hosts and HIV-1 infected people^23–26^, could be another reason for the enhanced pathogenicity of X4 tropic HIV-1. In nonprogressive SIV infection, a low viral burden in the CM CD4 subset and the lack of chronic immune activation are considered as two hallmarks^26–28^. While in most HIV-1 infected people, the level of CD4 activation could achieve a steady state^15^, the level of CD4 activation increased in 40700 after acute infection. Further efforts are needed to determine whether a broader range of CD4 subset infection could lead to an enhanced immune activation in HIV-1 infection.

The current study has several limitations. First, we only focused on a single participant. Second, because samples from the tissues such as lymph nodes were not available, the observations were based on cells from peripheral blood. Future research using animal model could lead to a better understanding of the immunopathogenesis of this X4 tropic T/F HIV-1.

## Figure Legends

**Extended Data Fig. 1.**
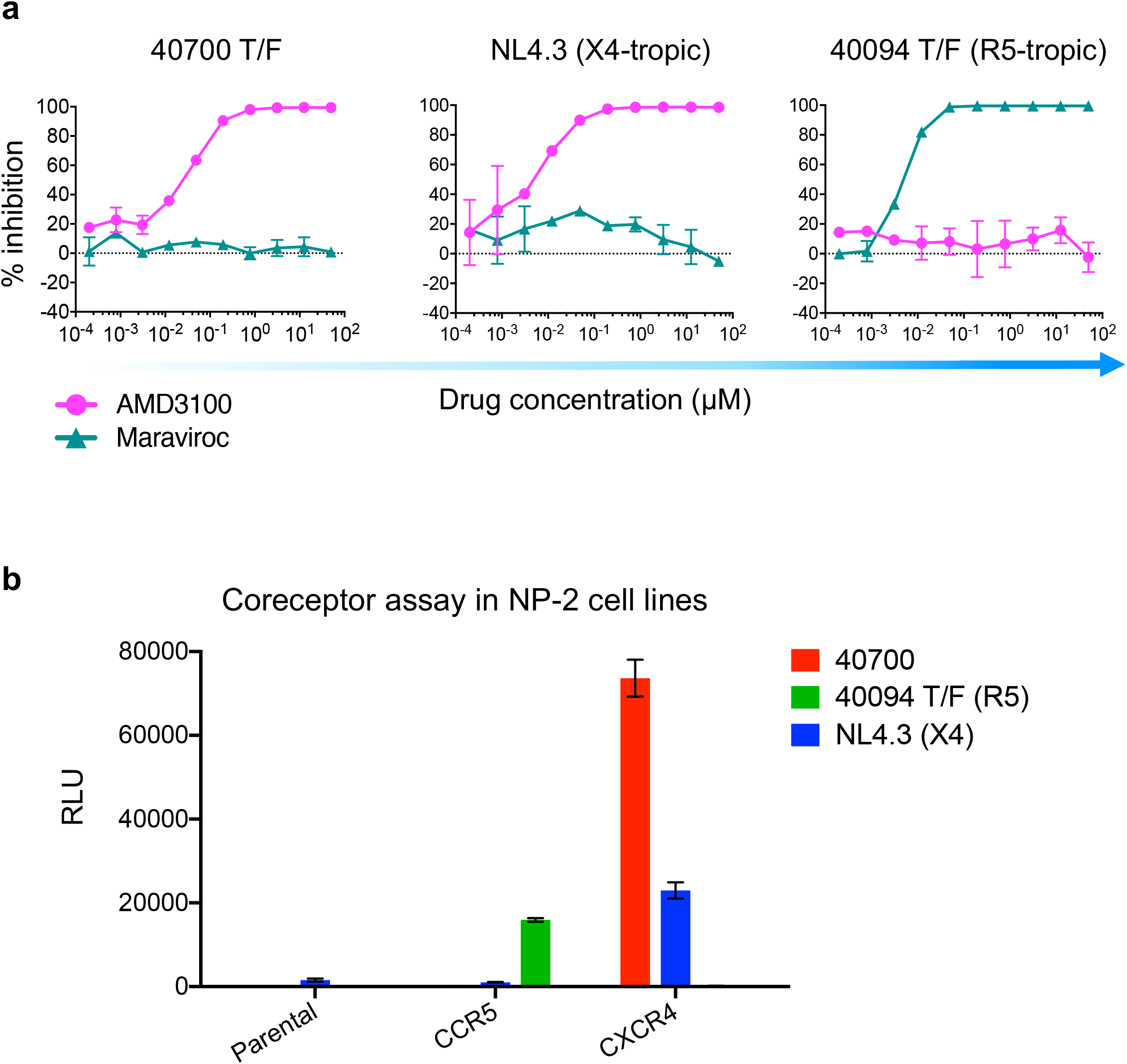
Determination of coreceptor usage of the 40700 T/F pseudovirus. **a**, TZM-bl cells were pre-treated with different concentrations of the CXCR4 inhibitor AMD3100 or the CCR5 inhibitor Maraviroc for 1 hour before infection. The X4 tropic virus NL4.3 and an R5 tropic T/F virus from participant 40094 were used as controls. The inhibition assays were performed in duplicate. The error bar shows the SD. **b**, Determination of coreceptor usage of the 40700 pseudovirus in NP-2 cell lines expressing CCR5 or CXCR4. Viral infectivity was considered positive if the RLU value was at least 5-fold higher than the background RLU value in the NP-2 parental cell line. The X4 tropic strain NL4.3 and an R5 tropic T/F virus from participant 40094 were used as controls. The experiments were performed in duplicate. The error bar represents the SD.

**Extended Data Fig. 2.**
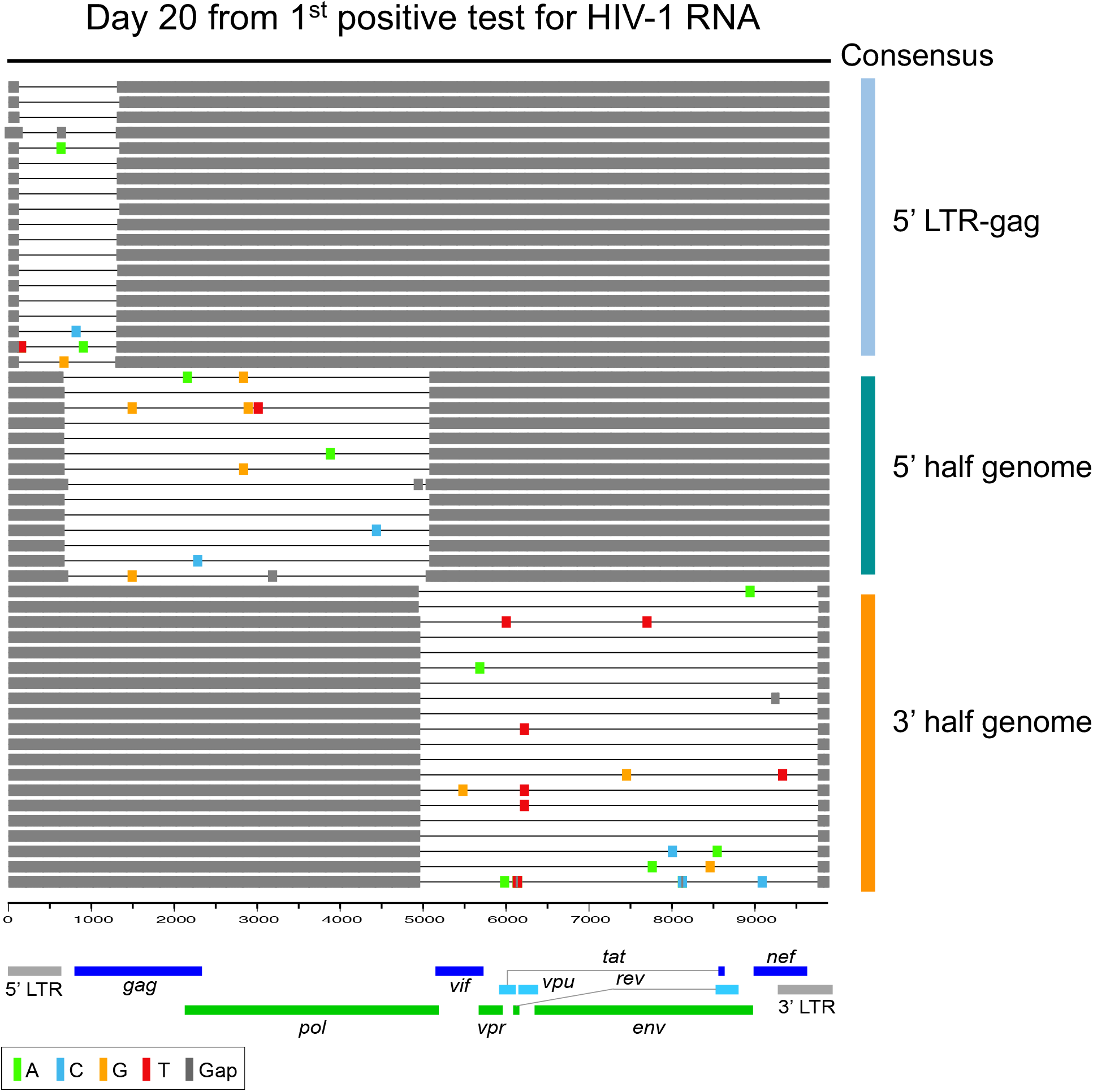
Inference of the full-length 40700 T/F viral genome. The highlighter plot shows alignment of SGA-derived viral sequences obtained at day 20 (from the 1^st^ positive test for HIV-1 RNA). Three overlapping fragments were amplified. The 5’ half and 3’ half viral genomes were amplified from plasma samples collected at day 20. The 5’ LTR-gag fragments were amplified from PBMCs collected at day 20. The black line on the top shows the consensus sequence of three overlapping fragments.

**Extended Data Fig. 3.**
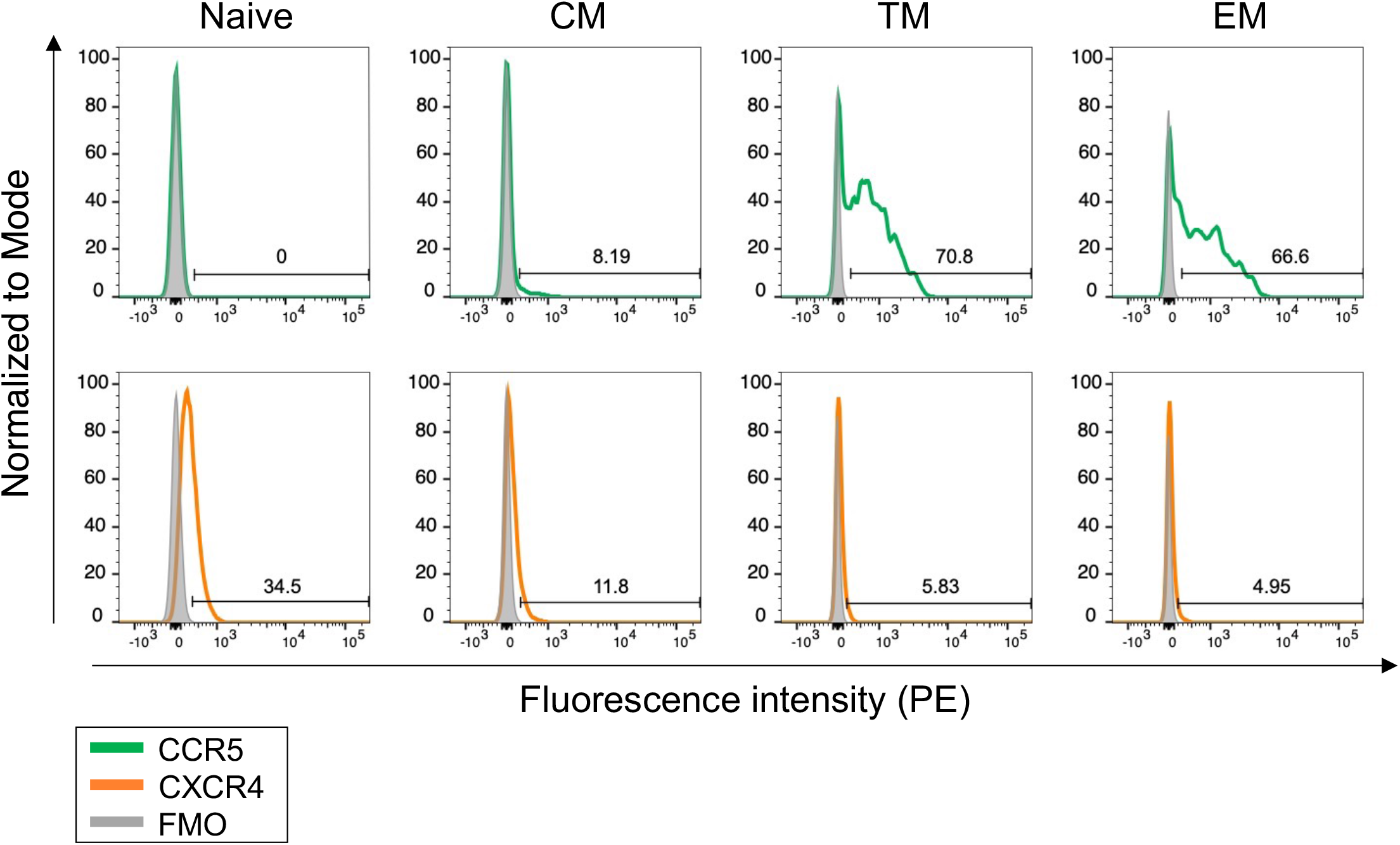
Determination of CCR5 and CXCR4 expression on CD4^+^ T cells from participant 40700. The expression of CCR5 and CXCR4 on each CD4 subset was determined using PBMCs collected at day 20 (from the 1^st^ positive test for HIV-1 RNA). The CCR5 and CXCR4 staining antibodies were titrated to determine the optimal concentrations. The flow data were collected using the DIVA 7.0 software on the FACSAria II (BD Biosciences) cell sorter/analyzer and analyzed by the FlowJo software. Positive cells are shown by percentage in each panel.

**Extended Data Fig. 4.**
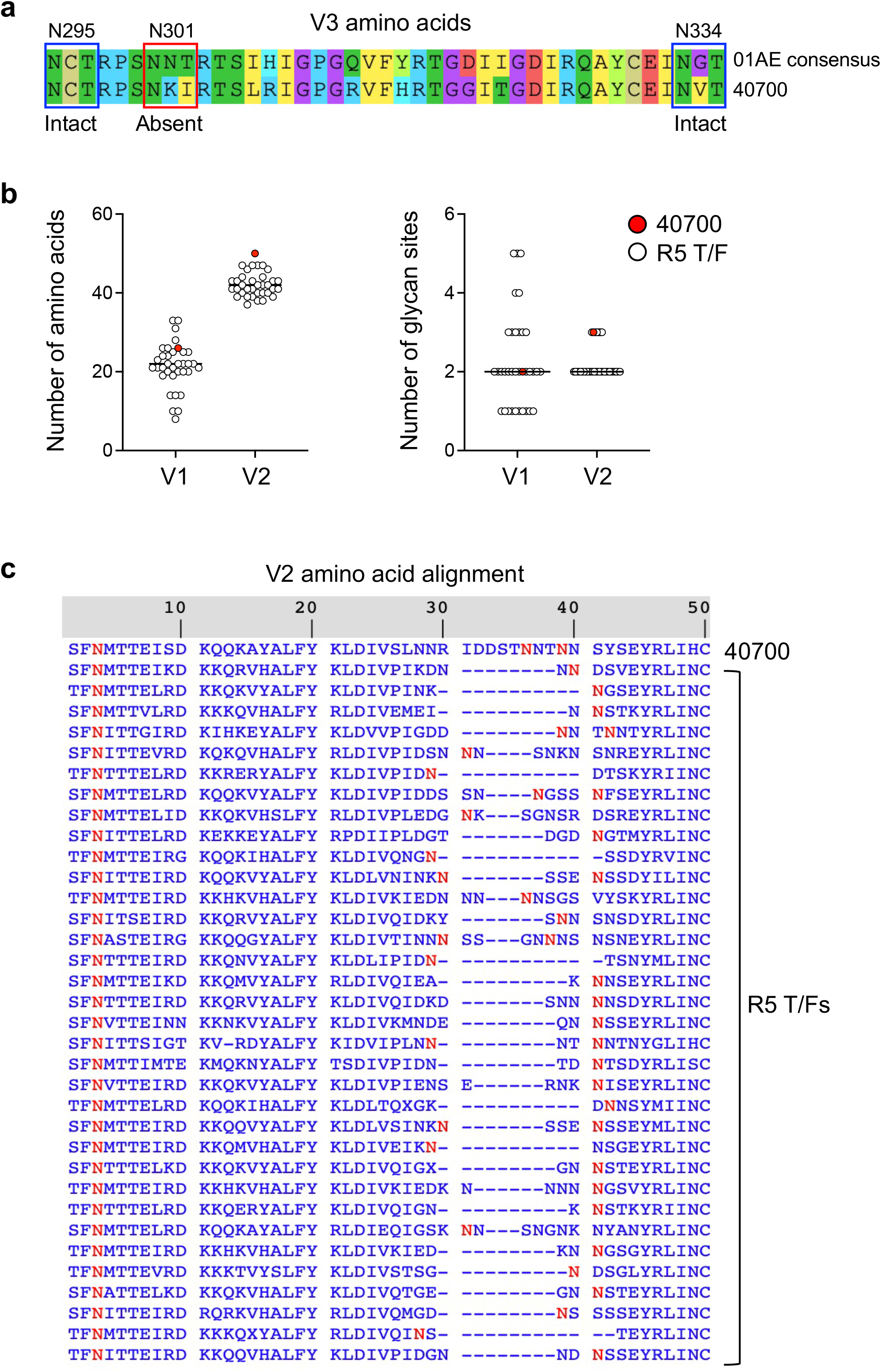
Amino acid sequences and glycan arrangement in V3 and V1/V2 regions of the 40700 T/F virus. **a**, The V3 amino acid sequence of the 40700 T/F virus compared with the CRF01_AE consensus sequence. Three conserved V3 glycan sites in CRF01_AE (N295, N301, and N334) are indicated by boxes. **b,** The length of the V1/V2 regions (left) and the number of V1/V2 glycan sites (right) in the 40700 T/F virus compared with R5 T/F viruses in the RV217 Thailand cohort. The 40700 T/F virus is shown in red. **c**, The V2 amino acid sequence and N-glycan sites in the 40700 T/F virus compared with R5 T/F viruses in the same cohort. The V2 amino acid sequences were obtained by Gene Cutter in the Los Alamos HIV Sequence Database. The N-glycan site (shown in red) was determined using the N-GlycoSite tool in the Los Alamos HIV Sequence Database.

## Methods

### Study participants

All study participants were from the RV217 Thailand acute infection cohort^9^. Participant 40700 became infected around the time of entry and the risk factor for HIV-1 transmission was men who have sex with men (MSM). The subtype of the 40700 T/F virus is CRF01_AE. Participant 40700 initiated ART on day 266 (from the 1^st^ positive test for HIV-1 RNA) and stopped ART on day 282. ART was restarted on day 388. Written consent was provided by all participants. The study was approved by the local ethics review boards, the Walter Reed Army Institute of Research, and the institutional review boards of the University of Maryland School of Medicine.

### Single genome amplification

Single genome amplification was carried out as previously described^29^. HIV-1 RNA in plasma was extracted using the QIAamp Viral RNA Mini Kit (Qiagen). To amplify the 3’ half viral genome, cDNA was synthesized by the SuperScript III reverse transcriptase (Invitrogen) using the primer 1.R3.B3R 5’-ACTACTTGAAGCACTCAAGGCAAGCTTTATTG-3’ (nt 9642-9611 in HXB2). The first round PCR was performed using the primers 07For7 5’-CAAATTAYAAAAATTCAAAATTTTCGGGTTTATTACAG-3’ (nt 4875-4912) and 2.R3.B6R 5’-TGAAGCACTCAAGGCAAGCTTTATTGAGGC-3’ (nt 9636–9607), and the second round PCR was performed with the primers VIF1 5’-GGGTTTATTACAGGGACAGCAGAG-3’ (nt 4900-4923) and Low2c 5’-TGAGGCTTAAGCAGTGGGTTCC-3’ (nt 9591-9612). To amplify the 5’ half viral genome, cDNA was synthesized using the primer 07Rev8 5’-CCTARTGGGATGTGTACTTCTGAACTT-3’ (nt 5193-5219). The first round PCR was performed using the primers 1.U5.B1F 5’-CCTTGAGTGCTTCAAGTAGTGTGTGCCCGTCTGT-3’ (nt 538-571) and 07Rev8 5’-CCTARTGGGATGTGTACTTCTGAACTT-3’ (nt 5193-5219), and the second round PCR was performed using the primers Upper1A 5’-AGTGGCGCCCGAACAGG-3’ (nt 634-650) and Rev11 5’-ATCATCACCTGCCATCTGTTTTCCAT-3’ (nt 5041-5066). Two microliters of the first round PCR products were used for the second round PCR amplification. The PCR conditions were as follows: one cycle at 94°C for 2 min; 35 cycles of a denaturing step at 94°C for 15 sec, an annealing step at 60°C for 30 sec, an extension step at 68°C for 4 min; and one cycle of an additional extension at 68°C for 10 min. The PCR amplicons were directly sequenced by the cycle sequencing and dye terminator methods. Individual sequences were assembled and edited using Sequencher (Gene Codes). The sequences were aligned using the Gene Cutter in the Los Alamos HIV Sequence Database followed by manual adjustment to obtain the optimal alignment.

### Pseudovirus preparation and titration

The pseudovirus stocks were prepared as previously described^30^. In brief, 2 μg of *env* clone was co-transfected with 4 μg of the pNL4.3-ΔEnv-vpr+-luc+ into 293T cells in a T25 flask using the FuGENE6 transfection reagent (Promega). The cells were cultured at 37°C for 6 hours before the medium was completely replaced with fresh medium. The culture supernatants containing the pseudoviruses were harvested at 72 hours post transfection, aliquoted and stored at −80°C until use. The infectious titers (TCID_50_) of the pseudovirus stocks were determined on TZM-bl cells.

### Generation of infectious molecular clone (IMC) of the 40700 T/F virus

To obtain the full-length genome of the 40700 T/F virus, three overlapping fragments were amplified using plasma or PBMC samples collected at day 20 (from the 1^st^ positive test for HIV-1 RNA) (Extended Data Fig. 2). The 5’ half and 3’ half viral genomes were amplified using day 20 plasma sample as described above. The 5’ LTR-gag fragment was amplified using viral DNA extracted from the day 20 PBMCs with the forward primer 5’-TGGAAGGGCTAATTTACTCCAAGAAAAG-3’ (nt 1-28) and the reverse primer 5’-TCTGATAATGCTGWRAACATGGGTAT-3’ (nt 1294-1319). The full-length 40700 T/F virus was inferred as the consensus sequence of the three overlapping fragments (Extended Data Fig. 2). The 40700 T/F sequence was chemically synthesized and cloned into the vector pUC57-Brick (Genescript).

To generate the viral stock of the 40700 T/F IMC, 6 μg of IMC was transfected into 293T cells in a T25 flask using the FuGENE6 transfection reagent (Promega). The cells were cultured at 37°C for 6 hours before the medium was replaced by fresh medium. The culture supernatants were harvested at 72 hours post transfection, aliquoted and stored at −80°C until use. The infectious titer (TCID_50_) of the viral stock was determined in TZM-bl cells.

### Determination of coreceptor usage

Coreceptor usage of the 40700 T/F pseudovirus was determined by both coreceptor inhibition assay in TZM-bl cell line and entry assay in NP-2 cell lines. For inhibition assay, TZM-bl cells were seeded in a 96-well plate at a density of 1 × 10^5^ cells per well. The next day, the cells were pre-treated with different concentrations of the CCR5 inhibitor Maraviroc or the CXCR4 inhibitor AMD3100 at 37°C for 1 hour. The treated cells were infected with approximately 500 TICD_50_ of the pseudovirus. The infected cells were lysed at day 3 after infection. The infectivity in each well was determined by measuring the relative luciferase units (RLU) in the cell lysates using the Britelite plus system (PerkinElmer). The percentage of inhibition was determined by comparing the infectivity with positive control wells without drug inhibition.

For entry assay, NP-2 cell lines expressing CCR5 or CXCR4 were seeded in a 96-well plate at a density of 1 × 10^5^ cells per well. The next day, the cells were infected with approximately 200 TCID_50_ of each pseudovirus (MOI = 0.002). After 6 hours of incubation at 37°C, the infected cells were washed twice with the culture medium and cultured at 37°C for three days. At 72 hours post infection, the infected cells were lysed, and the infectivity was determined by measuring the relative luciferase units (RLU) in the cell lysates using the Britelite plus system (PerkinElmer). Viral infectivity was considered positive if the RLU value was at least 5-fold higher than the background RLU value in the NP-2 parental cell line. All experiments were performed in triplicate.

Coreceptor usage of the 40700 IMC was determined in NP-2 cell lines expressing CCR5, CXCR4 and a panel of alternative coreceptors (CCR3, APJ, FPRL1, CCR8, GPR15, CCR2b and CCR1). NP-2 cells were seeded in a 96-well plate one day before infection at a density of 1 × 10^5^ cells per well. Approximately 100 TCID_50_ of viral stock was used for infection (MOI = 0.001). After 4 hours of incubation at 37°C, the infected cells were washed three times with the culture medium and cultured at 37°C for five days. The p24 concentrations in the culture supernatants were measured on day 5 post infection (PerkinElmer).

### Determination of viral replication capacity in primary CD4^+^ T cells

Purified CD4^+^ T cells from healthy donors were stimulated with 1 µg/mL soluble anti-CD3 (clone OKT3, eBioscience) and 1 µg/mL soluble anti-CD28 (clone CD28.2, eBioscience) in the presence of 50 IU/mL IL-2 (PeproTech) for three days. The stimulated CD4^+^ T cells were infected by the 40700 T/F IMC at 37°C for 4 hours (MOI = 0.001). The cells were washed three times after infection and were cultured at 37°C for 7 days. The culture supernatants were collected every day and the viral growth kinetics was determined by measuring the p24 concentration in the supernatant (PerkinElmer). All infections were performed in triplicate.

### CCR5 genotyping

Genomic DNA (gDNA) was extracted from 1×10^6^ PBMCs using the QIAamp DNA Blood Mini Kit (Qiagen) per manufacturers’ instructions. To differentiate between CCR5wt/wt, Δ32/Δ32, and wt/Δ32 genotypes, two primer amplification strategies were employed for each sample. Forward primers CCR5delF1 (5’-ACCGTCAGTATCAATTCTGGAAGA-3’) and CCR5span1a (5’-CATTTTCCATACATTAAAGATAGT-3’) were used to specifically detect the CCR5wt and CCR5Δ32 genotypes, respectively; the reverse primer CCR5del2 (5’-CATGATGGTGAAGATAAGCCTCACA-3’) was common for both (all Sigma-Aldrich; St. Louis, MO). Two PCR master mixes differing only in the forward primer were prepared at a final volume of 10 µL, including 5 µL Platinum SYBR Green qPCR Supermix-UDG with ROX (Invitrogen), 0.1 µM CCR5del2, and 0.1 µM CCR5span1a or CCR5delF1 primers and 10 ng of gDNA sample. PCR was performed in 384 well plates sealed with optical adhesive film (ABI), and amplification performed on Veriti thermal cyclers (ABI) using the following parameters: 50°C, 2 min; 95°C, 2 min; 40 cycles of 95°C, 15 sec and 61°C, 30 sec; 4°C. Dissociation curve analyses were then performed on the 7900HT Fast Real-Time PCR instrument using the SDS2.4 software to detect presence or absence of amplified products (both ABI).

### CD4 subset analysis and sorting

CD4 subset analysis and sorting were performed as described previously^12^. PBMC samples were stained by the following antibodies: CD3-Brilliant Violet 605 (clone OKT3, BioLegend), CD4-PerCP-Cy5.5 (clone OKT4, Biolegend), CCR7-PE-CF594 (clone 2-L1-A, BD Biosciences), CD27-PE (clone M-T271, BioLegend), CD45RO-APC (clone UCHL1, BioLegend). The cells were then stained by the Live-dead aqua (Invitrogen) prior to flow analysis. The stained cells were sorted on a BD FACSAria II cell sorter (BD Biosciences). Four CD4 subsets were defined as follows: naïve (CD45RO^-^, CCR7^+^, and CD27^+^), central memory (CD45RO^+^, CCR7^+^, and CD27^+^), transitional memory (CD45RO^+^, CCR7^-^, and CD27^+^) and effector memory (CD45RO^+^, CCR7^-^, and CD27^-^). The purity of each sorted CD4 subset was higher than 95%.

### Flow cytometry

To determine CCR5 and CXCR4 expression on each CD4 subset, PBMCs were stained with the following antibodies: CD3-Brilliant Violet 605 (clone OKT3, BioLegend), CD4-PerCP-Cy5.5 (clone OKT4, BioLegend), CCR7-PE-CF594 (clone 2-L1-A, BD Biosciences), CD27-FITC (clone M-T271, BioLegend), CD45RO-APC (clone UCHL1, BioLegend), CCR5-PE (clone J418F1, BioLegend) or CXCR4-PE (clone Q18A64, BioLegend). The CCR5 and CXCR4 antibodies were titrated to determine the optimal concentration. To determine the non-specific staining, cells was cold-inhibited by a 100-fold excess of the unlabeled CCR5 or CXCR4 antibody (the same clone as the staining antibody) mixed with the respective labeled antibody. A fluorescence minus one (FMO) staining was also determined for CCR5/CXCR4 staining. The highest concentration of the labeled antibody with which the cold inhibition showed virtually overlapping staining with the FMO was used to determine the levels of CCR5 and CXCR4 expression on each CD4 subset.

To quantify the expression of cell activation and proliferation markers, PBMCs were stained with the following antibodies: CD3-Brilliant Violet 605 (clone OKT3, BioLegend), CD4-Alexa Fluor 700 (clone OKT4, BioLegend), CCR7-BV421 (clone G043H7, BioLegend), CD27-PerCP-Cy5.5 (clone M-T271, BioLegend), CD45RO-APC/Fire 750 (clone UCHL1, BioLegend), Ki-67-APC (clone Ki-67, BioLegend), HLA-DR-PerCP (clone L243, BioLegend), CD38-PE-594 (clone HIT2, BioLegend), and PD1-PE (clone EH12.2H7, BioLegend). The flow data were collected using the DIVA 7.0 software on the FACSAria II (BD Biosciences) cell sorter/analyzer and analyzed by the FlowJo software (FlowJo LLC, Ashland, OR).

### Quantification of total and integrated HIV-1 DNA

Total and integrated HIV-1 DNA were quantified as previously described^31^. The exact number of cells in each PCR reaction was determined by amplifying the human CD3 gene using the primers HCD3OUT5 5’-ACTGACATGGAACAGGGGAAG-3’ and HCD3OUT3 5’-CCAGCTCTGAAGTAGGGAACATAT-3’. To quantify the total HIV-1 DNA, the first round PCR was performed using the primers ULF1 5’-ATGCCACGTAAGCGAAACTCTGGGTCTCTCTDGTTAGAC-3’ (nt 436-471, 9521-9556), UR1 5’-CCATCTCTCTCCTTCTAGC-3’ (nt 775-793), HCD3OUT5 and HCD3OUT3. To quantify integrated HIV-1 DNA, the first round PCR was performed using the primers ULF1, Alu1 5’-TCCCAGCTACTGGGGAGGCTGAGG-3’ (nt 8674-8697), Alu2 5’-GCCTCCCAAAGTGCTGGGATTACAG-3’ (nt 6237-6261), HCD3OUT5 and HCD3OUT3. The first-round PCR conditions were as follows: a denaturation step at 95°C for 8 min; 12 cycles of a denaturing step at 95°C for 1 min, an annealing step at 55°C for 40 sec (1 min for integrated DNA), an extension step at 72°C for 1 min (10 min for integrated DNA), followed by an elongation step at 72°C for 15 min. For all experiments, the first round PCR products were diluted 10-fold and a total of 6.4 µL of the diluted PCR products were used for real-time PCR on the QuantStudio 3 Real-Time PCR Systems (Thermo Fisher Scientific) using the Perfecta qPCR ToughMix (QuantaBio). The copy number of the total and integrated HIV-1 DNA were quantified using the primers Lambda T 5’-ATGCCACGTAAGCGAAACT-3’ (nt 1555-1572), UR2 5’-CTGAGGGATCTCTAGTTACC-3’ (nt 583-602, 9668-9687), and the probe UHIV TaqMan 5’-/56-FAM/CACTCAAGG/ZEN/CAAGCTTTATTGAGGC/3IABkFQ/-3’ (nt 522-546, 9607-9631). The copy numbers of human CD3 gene were determined using the primers HCD3IN5’ 5’-GGCTATCATTCTTCTTCAAGGT-3’, HCD3IN3’ 5’-CCTCTCTTCAGCCATTTAAGTA-3’, and the probe CD3 TaqMan 5’-/56-FAM/AGCAGAGAA/ZEN/CAGTTAAGAGCCTCCAT/3IABkFQ/-3’. The real-time PCR conditions were as follows: A denaturing step at 95°C for 4 min, 40 cycles of a denaturing step at 95°C for 3 sec, an annealing and extension step at 60°C for 20 sec. The ACH2 cell lysates were used for standard curve.

### Quantification of cell associated HIV-1 RNA

To quantify cell-associated HIV-1 RNA, RNA was extracted from sorted CD4^+^ T cells using the RNeasy Mini kit (Qiagen). A total of 8.5 µL extracted RNA was subjected to one-step RT-PCR using the Superscript III one-step RT-PCR system (Invitrogen). To amplify the *pol* region, the one-step RT-PCR was performed using the forward primer Pol F1 5’-TACAGTGCAGGGGAAAGAATA-3’ (nt 4809-4829) and the reverse primer Pol R1 5’-CTTCTTGGCACTACTTTTATGTCAC-3’ (nt 4993-5017). The PCR conditions were as follow: a reverse transcription step at 50°C for 1h; A denaturing step at 94°C for 2 min; 16 cycles of a denaturing step at 94°C for 15 sec, an annealing step at 55°C for 30 sec, an extension step at 68°C for 1 min, and one cycle of an additional extension at 68°C for 5 min. The first round PCR products were diluted 10-fold and a total of 6.4 µL of diluted PCR products were used for the real-time PCR using the forward primer Pol F1, the reverse primer Pol R2 5’-CTGCCCCTTCACCTTTCC-3’ (nt 4957-4974), and the probe Pol Famzen: 5’-/56-FAM/TTTCGGGTT/ZEN/TATTACAGGGACAGCAG/3IABkFQ/-3’ (nt 4896-4921). The real-time PCR was performed on the QuantStudio 3 Real-Time PCR Systems (Thermo Fisher Scientific) using the following conditions: A denaturing step at 94°C for 4 min, 45 cycles of a denaturing step at 94°C for 3 sec, an annealing and extension step at 60°C for 20 sec. To amplify the *tat/rev* transcript, the one-step RT-PCR was performed using the forward primer Tat1.4 AE 5’-TGGCAG GAAGAAGCGGAAG (nt 5971-5989) and the reverse primer Rev AE-ter 5’-TGTCTCTGYCTTGCTCKCCACC-3’ (nt 8433-8454). The real-time PCR was performed using the forward primer Tat2 AE-bis 5’-GTAAGGATCATCAAAATCCTVTACCARAGCA-3’ (nt 6015-6045), the reverse primer Rev AE-ter 5’-TGTCTCTGYCTTGCTCKCCACC-3’ (nt 8433-8454), and the probe Tat-Rev AE 5’-/56-FAM/TT CYT TCG G/ZEN/G CCT GTC GGG TTC C/3IABkFQ/-3’ (nt 8399-8421). The PCR conditions were the same as for amplifying the *pol* region. The copy number of the input RNA was determined by using the RNA standard generated by *in vitro* transcription. In brief, the amplicon region (the CRF01_AE consensus sequence was used in the current study) was cloned into the pUC57 vector downstream of the T7 promoter. The DNA fragment containing the amplicon was PCR amplified and the RNA was generated by *in vitro* transcription using the MEGAscript T7 Transcription Kit (Invitrogen).

### Neutralization assay

The neutralization activity of plasma samples and monoclonal antibodies (mAbs) was determined by using a luciferase reporter system in TZM-bl cells. Plasma samples were heat inactivated at 56°C for 45 minutes. The inactivated plasma was diluted at a 1:3 serial dilution starting from 1:20. The mAbs were diluted at a 1:3 serial dilution from a starting concentration of 25 μg/mL. The virus stocks were diluted to a concentration that achieved approximately 150,000 RLU in the TZM-bl cells (or at least 10 times above the background RLU of the cells control). The serial diluted plasma samples or mAbs were then incubated with the viruses for 1 hour at 37°C in duplicate before the TZM-bl cells were added. The 50% inhibitory dose (ID_50_) was determined as the dilution at which the relative luminescence units (RLUs) were reduced by 50% in comparison to the RLUs in the virus control wells after subtraction of the background RLUs in cell control wells.

### Determination of the rate of CD4^+^ T cell decline

The rate of CD4 decline was determined using a linear mixed effect model (LME). The LME model was hierarchical in the sense that it estimated a population specific slope and intercept with time, as well as subject-specific slopes and intercepts. The longitudinal CD4 data from the earliest available time point to the last available time point before ART initiation was used for analysis. To determine whether participant 40700 had a statistically significant rate of CD4 decline, we calculated the empirical best linear unbiased prediction (EBLUP) for the rate of CD4 decline. A normal distribution test was then conducted to determine the statistical significance.

## Acknowledgements

We thank the study participants of the RV217 Thailand cohort. This study was supported by the Institute of Human Virology, University of Maryland School of Medicine. Part of the study was supported by cooperative agreements between the Henry M. Jackson Foundation for the Advancement of Military Medicine, Inc., and the U.S. Department of Defense (DOD). M.H.M. and H.S. were supported by the NIH grant R21AI147893.

## Disclaimer

The views expressed are those of the authors and should not be construed to represent the positions of the U.S. Army, the Department of Defense, the National Institutes of Health, the Department of Health and Human Services, or the Henry M. Jackson Foundation for the Advancement of Military Medicine, Inc. The investigators have adhered to the policies for protection of human subjects as prescribed in AR-70-25.

## Competing interests

The authors declare no competing interests.

## Data availability

All newly generated nucleic acid sequences in the current study were deposited in GenBank with accession numbers OR231807-OR231874.

## Notes

### Competing Interest Statement

The authors have declared no competing interest.

